# Large-scale two-photon imaging revealed super-sparse population codes in V1 superficial layer of awake monkeys

**DOI:** 10.1101/252940

**Authors:** Shiming Tang, Yimeng Zhang, Zhihao Li, Ming Li, Fang Liu, Hongfei Jiang, Tai Sing Lee

## Abstract

Efficient coding has been proposed as a general principle for the sensory systems. The efficient coding hypothesis predicts that neuronal population responses should be sparse, but limited by the measurement techniques, the precise estimates of the population sparseness of visual cortical neurons are still uncertain. Here, we employed large-scale two-photon calcium imaging to examine the neuronal population activities in V1 superficial layers of awake macaques in response to a large set of natural images. We found that only 0.5% of these neurons on average responded strongly to any given natural image with response strength above half of their individual peak responses, which is more than tenfold sparse over those reported by early studies. We further showed that these sparse population activities contain sufficient information for discriminating images with high accuracy. This study provided the first accurate measure of sparseness in V1 neuronal population responses, which support super-sparse neural codes in primates.

## Introduction

Sparse coding is an important organizing principle in the sensory system (Barlow, 1981; Olshausen and Field, 1996). The efficient coding hypothesis predicts that neuronal population responses should be sparse, though the optimal level of sparseness depends on many factors. Most of the experimental evidences in support of sparse coding is based on the sparseness of an individual neuron’s responses to natural images, measured using single-unit recording techniques (Haider et al., 2010; Hromadka et al., 2008; Rust and DiCarlo, 2012; Vinje and Gallant, 2000). One direct measurement of population sparseness was provided by a two-photon imaging (GCaMP6f) study on rodents (Froudarakis et al., 2014). The absolute level of estimated sparseness depends on multiple factors. However, the saturation (above 60-80Hz) in GCaMP6 signal can affect the estimate of sparseness (Chen et al., 2013; Froudarakis et al., 2014). Thus, the precise measurement of population sparseness of neuronal response, particularly in primates, remains lacking.

In this study, we provided the first measurement of population sparseness of primate V1, by using two-photon calcium imaging on a large population of neurons at single cell resolution. We used calcium indicator GCaMP5 derived from AAVs (Akerboom et al., 2012), which we have shown earlier to exhibit linear behavior across a much wider range of firing rates (10 Hz to 150 Hz) (Li et al., 2017). This allows us to measure more accurately the response sparseness of almost an entire population of densely packed neurons in superficial layers 2/3 of V1 in an 850 μm × 850 μm field of view - the scale of about 1-2 hypercolumn.

## Results and discussion

We employed two-photon imaging with GCaMP5 to measure the neuronal population responses (about 1000 neurons per monkey) in V1 layers 2/3 of two awake macaques to 2,250 natural images. The neurons’ calcium signals in response to visual stimuli were recorded while the monkey performed a fixation task. During each fixation trial, a blank screen was presented for one second, followed by a visual stimulus for another one second. The ROI of an activated cell was identified when the brightness of a compact region (>25 pixels) exceeded 3 stds (standard deviations) in each single differential image. The ratio of fluorescence change (Δ*F/F0*) of these ROIs was calculated for each activated cell. *ΔF = F‐ F0*, where *F0* is the baseline activity during the blank screen prior stimulus onset in each trial and *F* is fluorescence activity in the ROI during stimulus presentation in the trial. A neuropil-correction was performed with an index of 0.7 (Chen et al., 2013).

The receptive fields (RFs) of the neurons were first localized using oriented gratings and bars presented in different positions. The RF centers of imaged neurons were located between 3^o^ and 5^o^ in eccentricity. In each trial, a stimulus of size 4^o^ × 4^o^ randomly drawn from a set of 2,250 natural image stimuli (Figure 1, Figure 1-Figure supplements 1c) was presented. The entire set of stimuli was repeated for three times. These natural images evoked robust visual responses in the imaged neurons (Figure 1a,b, Figure 1-Figure supplements 1c and 2).

**Figure 1.**
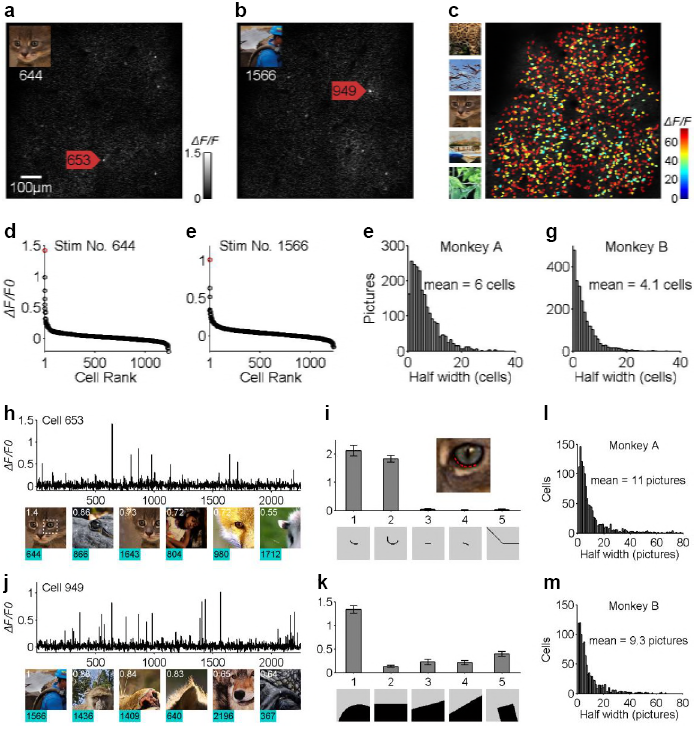
Sparse neuronal responses of V1 layer 2/3 neurons to natural scenes. (a and b) Ca images of the neuronal population response to two different natural images as shown in the insets. Typically, only a few neurons, among the nearly 1000 neurons measured (1,225 neurons for Monkey A or 982 neurons for Monkey B), responded strongly to a single patch of natural scenes. (c) The overall neuronal population responses to all 2,250 natural images. Eachcellwascolor-codedaccording tothe response intensity toits optimal stimulus respectively. (d and e) The distributions of neuronal population responses to the two natural images respectively. Abscissa indicates the 1,225 neurons that showed significant response to natural images, in ranked order according to their responses to each image. Ordinate indicates Δ*F/F0*. Cells 653 and 949 are colored red respectively. (f and g) The distribution of the sharpness measures of population response distribution in terms of half-height width across the natural stimuli tested for the two monkeys. Less than 0.5% of the cells responded above half of their individual peak response to any given natural image. (h and i) The response of one example cell (cell 653) to the entire set of natural scene stimuli, exhibiting high stimulus specificity. (j and k) Another example cell (cell 949) also shows high stimulus specificity. (l and m) The distributions of stimulus specificity of neurons, in terms of half-height width of the stimulus tuning curves. Each cell would typically respond strongly to less than 0.5% of the natural images in our test set.

We examined the neuronal population responses to the natural image set. We found that out of the 1,225 neurons in monkey A and 982 in monkey B, only a few neurons from each monkey strongly responded to each image patch (Figure 1a,b), though a large number of imaged neurons could be activated by the whole stimulus set (Figure 1c). The rank-ordered distributions of the population responses were always sharply peaked (Figure 1d,e). On average, the half-height bandwidths of population response distributions were 0.49% (6.0/1,225) and 0.42% (4.1/982) for monkey A and B respectively (Figure 1f,g). In other words, only about 0.5% of the cells responded substantially, each with activity level above half of its observed peak response, indicating a very high degree of sparseness in population responses (***see Methods***).

We also examined each neuron’s stimulus specificity, or its life-time sparseness. We found most cells responded substantially only to a small number of images in the whole stimulus set (Figure 1h,j). Interestingly, the preferred images for individual neuron often shared common features. For example, neuron 653 of monkey A was most excited when its receptive field (0.8^o^ in diameter) covered the lower rim of the cat’s eye (indicated by the red dashed line in the inset in Figure 1i). The neuron’s preference for the specific curve feature was further confirmed by a subsequent experiment testing the neurons’ selectivity to many artificial patterns (Figure 1i; see also Tang et al., 2017). Similarly, neuron 949 of Monkey A was found to be selective to an opposite curvature embedded in its preferred natural stimulus set (Figure 1k). These observations are consistent with earlier observations that V1 neurons might be selective to more complex patterns (Hedgé and Van Essen, 2007; Tang et al., 2017), which would result in highly sparse population responses. The average half-height stimulus bandwidths in stimulus tuning were 0.49% (11/2,250) and 0.41% (9.3/2,250) for monkeys A and B (Figure 1l,m) respectively. This high degree of stimulus specificity or life-time sparseness goes hand in hand with the high degree of measured population sparseness.

To understand how much information was carried in the sparse ensemble of population activities, we evaluated how well the sparse neural responses allow a decoder to discriminate the 2,250 stimuli (Quiroga RQ and Panzeri, 2009; Froudarakis et al., 2014). When the entire population activities were included, the decoding accuracies were 54% for monkey A and 38% for monkey B, whereas the chance accuracy was only 0.04% (Figure 2a,b). In the following, we refer to these decoding accuracies as “achievable decoding accuracies” for subsequent comparison.

**Figure 2.**
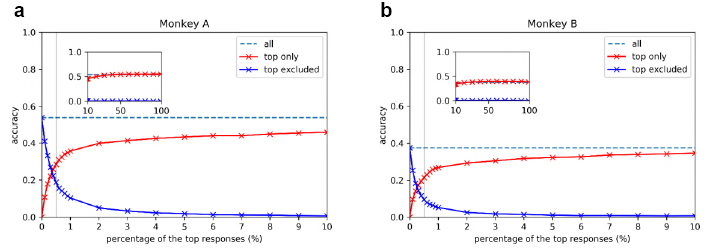
Performance on the image decoding task using strong and weak neural responses. Y axes show the cross-validated decoding accuracy on the 2250-way image classification task, and X axes show the percentage of top responses included (red curves) and excluded (blue curves). Dash lines are the referential “achievable decoding accuracies” using all the neural responses. Gray vertical lines highlight the decoding accuracies including or excluding the top 0.5% responses.

To examine how much information the downstream neurons could derive from the strongest 0.5% responses, we set all the responses below the top 0.5% to 0 and used only the top 0.5% strong responses to perform decoding (top only). Remarkably, a decoding accuracy of 28% could be achieved for monkey A and 21% for monkey B, which were about 50% of achievable decoding accuracies for both monkeys.

We also studied how well the decoding could do when the top 0.5% sparse responses were excluded by setting them to 0 (top excluded). We found by excluding the top 0.5% top responses, over 50% of the achievable accuracy was lost for each monkey. Thus, we showed that these strong and sparse responses convey both the necessary and sufficient information to downstream neurons for decoding the images with high accuracy.

The decoding accuracies under different percentages of the top responses included (red curve) or excluded (blue curve) revealed that most of the achievable decoding accuracy was contributed by the top 5% of the strong responses in this 2,250-way decoding task (Fig. 2).

Thus, our study provided the first direct measurement of sparseness of large-scale neuronal population responses in V1 superficial layer of awake macaque monkeys, using large-scale two-photon imaging. We found that a very small ensemble of neurons in V1 superficial layer would be active concurrently in response to any given natural image. Using decoding analysis, we showed that these small ensembles of neural responses provide necessary and sufficient information for downstream neurons for image discrimination.

Earlier studies inferred population sparseness based on measurement of life-time sparseness. We showed that population sparseness measure was indeed comparable to life-time sparseness measure. However, the level of sparsity we observed (0.5% at half height) was considerably higher than earlier life-time sparseness estimates based on single unit recording in macaque (Rust and DiCarlo, 2012). Studies on rodents yielded a considerable range of estimates on response sparseness which varied with measuring techniques (Haider et al., 2010; Hrimadka et al., 2008; Froudarakis et al., 2014). The single-unit study using the cell-attached technique might be the most accurate (Hromadka et al., 2008), which showed that neurons were mostly silent in the awake auditory cortex, and inferred that less than 5% of the neurons would be strongly active (above 20 Hz) at each given instance. This sparse measure is actually consistent with our data (bandwidth at about 25% height maximum), which confirmed that most of early study on neuronal sparseness with traditional extracellular recording were debatable from technical angle. From our observation, there were densely packed neurons with small cell bodies in the superficial layers of V1. It will be hard to get well isolated and long-term stable single-unit signals with extracellular recording method. Our study have eliminated the bias in neuron sampling and probed the neurons with large set of natural stimuli, thus provide a direct sparseness measurement of V1 neurons in primates at both a population and single-cell level.

The high level of sparsenes showever is consistent with two recent interesting conjectures in theoretical neuroscience. First, based on metabolic reasons and the cost of spiking, it has been conjectured that fewer than 1% of the neurons should be substantially active concurrently in any brain area (***Lennin, 2003***). Second, and more importantly, theoretical sparse coding studies have suggested that because the number of V1 neurons is at least 200 times more abundant than its thalamic input, V1 neurons could be quite specialized in their feature tuning and highly sparse in their population responses (Olshausen, 2013; Rehn and Sommer, 2007), as we have observed using two-photon imaging techniques in the V1 superficial layer (Tang et al. 2017). The observed high degree of sparseness and complexity of V1 superficial-layer neurons have consequential implications on our understanding of efficient coding in the brain.

## Materials and methods

All experimental protocols were approved by the Peking University Animal Care and Use Committee.

### Subjects

The study used four adult rhesus monkeys (A and B), 4–5 years of age and weighing 5–7 kg. Two sequential surgeries were performed on each animal under general anesthesia and strictly sterile conditions. In the first surgery, a 16-mm hole was drilled in the skull over V1. The dura was opened to explore the cortex, into which 50-100 nl AAV1.hSynap.GCaMP5G.WPRE.SV40 (AV-1-PV2478, titer 2.37e13 (GC/ml), Penn Vector Core) was pressure-injected at a depth of ∼500 μm. After AAV injection, the dura was sutured, the skull cap was placed back, and the scalp was sutured. Then the animal was returned to its cage for recovery. Antibiotic (Ceftriaxone sodium, Youcare Pharmaceutical Group Co. Ltd., China) was administered for one week. After 45 days, a second surgery was performed, in which three head-posts were implanted on each animal’s skull, two on the forehead and one on the back of the head. A T-shaped steel frame was connected to these head-posts for head stabilization during imaging. The skull and dura were later on opened again to explore the cortex. A glass cover-slip (diameter 8 mm and thickness 0.17 mm) was glued to a titanium ring, and then gently pushed down onto the cortical surface. A ring-shape GORE membrane (20 mm in outer diameter) was inserted under the dura. The titanium ring was glued to the dura and skull with dental acrylic to form an imaging chamber. The whole chamber (formed by thick dental acrylic) was covered by a steel shell to prevent breakage of the cover-slip when the animal was returned to the home cage.

### Behavioral task

During imaging, each monkey was seated in a primate chair with head restraint and performed a fixation task, which involved fixating on a small white spot (0.1°) within a window of 1° for over 2 seconds to obtain a juice reward. Eye position was monitored with an infrared eye-tracking system (ISCAN, Inc.) at 120 Hz.

### Visual stimuli

Visual stimuli were generated using the ViSaGe system (Cambridge Research Systems) and displayed on a 17” LCD monitor (Acer V173, 80Hz refresh rate) positioned 45 cm from the animal’s eyes. Each stimulus was presented for 1 second after a 1 second blank within a fixation period of 2 seconds. We estimated the RF sizes and positions of the imaged neurons with small drifting gratings and bars presented at different locations. The RFs were estimated to be 0.2° to 0.8° in size with RF locations between 3 to 5 degrees in eccentricity for both monkeys.

Drifting and oriented gratings were tested to examine the visual responses of imaged neurons. Small patches (0.8° in diameter) of gratings with 100% contrast square waves were presented to the center of RFs of imaged cells, with 2 spatial frequencies (4.0 and 8.0 cyc/deg) at 2 temporal frequencies (1 and 2 Hz), 6 orientations, and 2 directions (30° apart).

A natural stimulus set (NS) of 2,250 4° × 4° stimulus patches extracted from different natural scene photos was used to examine the neuronal responses to natural stimuli. The order of the stimuli was randomized in each session. These stimuli were tested on monkeys A and B, each with at least three repetitions.

### Eye movement control

We analyzed the distribution of eye-positions during stimulus ON periods. The monkeys’ fixation during stimulus presentation (from 1 to 2 second in the graph) was quite stable and accurate. The distribution of eye positions during stimulus presentation, with standard deviations less than 0.05°, which was significantly smaller than the typical receptive field sizes of neurons at 3-5 degree eccentricities (range from 0.2 to 0.8 degrees). To examine whether the eye movement had significant contribution to the distribution of neuronal population responses, we compared the standard divisions (stds) of eye position in different neuronal population response cases. We considered three classes of population responses: (1) weak response (Δ*F/F0* < 0.5), (2) sparse strong response (one or two cells responded), (3) dense response (more than ten cells responded). We found no statistically significant differences in the distribution of eye position data in these three classes (Tang et al., 2017), which indicated the observed effects are not caused by movement differences. The ROC and decoding analysis (Figure 1-Figure supplement 2), demonstrating the reliability of the neural responses across trials, confirm that the sparse population responses were evoked by stimuli repeatedly, not by random eye-movement jitters.

### Two-photon imaging

After a recovery period of 10 days from the second surgery, the animals were trained to maintain eye-fixation. Two-photon imaging was performed using a Prairie Ultima IV (In Vivo) two-photon microscope (Bruker Nano, Inc., FMBU, formerly Prairie Technologies) and a Ti: Sapphire laser (Mai Tai eHP, Spectra Physics). The wavelength of the laser was set at 1000 nm. With a 16× objective (0.8-N.A., Nikon), an area of 850μm × 850μm was imaged. A standard slow galvo scanner was used to obtain static images of cells with high resolution (1024 × 1024). The fast and resonant scan (up to 32 frames per second) was used to obtain images of neuron activity. The images were recorded at 8 frames per second by averaging each 4 frames. The infected cells up to 700 μm in depth could be imaged. We mainly focused at the layer at 160 μm to 180 μm depth which contained a high density of infected cells.

### Imaging data analysis

All data analyses were performed using customized Matlab software (The MathWorks, Natick, MA). The images from each session were first realigned to a template image (the average image of 1000 frames) using a normalized cross-correlation-based translation algorithm, to correct the X-Y offset of images caused by the motion between the objective and the cortex.

The cell density was high in superficial V1, and many cell bodies were quite dim at rest. It was difficult to identify these cells directly by eye or by algorithm based on the morphology from their static images. Hence, we identified ROIs for cell bodies based on their responses. The differential images (average frame of the stimulus ON period subtracting that of stimulus OFF period for each stimulus condition) were first filtered using low-pass and high-pass Gaussian filters (5 pixels and 50 pixels, 2 orders respectively). Notably, these two filters were only used for ROI identifications. In all further analyses, we used the raw data without any filtering. Connected subsets of pixels (>25 pixels) with average pixel value greater than 3 standard deviations (std) in these differential images were identified as active neurons (ROIs). Note that this 3 std empirical value was used only for deciding the ROIs of activated cells and was not used as a cutoff threshold for measuring neuronal responses. We found that a higher std would allow the detection and selection of the ROIs of cell bodies more accurately but would miss some weakly responding cells, and a lower std threshold may include more cells but would have a greater chance of including some false ROIs that cannot be matched to any cell bodies. The ratio of fluorescence change *(ΔF/FO)* of these ROIs was calculated for each activated cell. ***Δ****F = F‐ F0,* where *F0* is the baseline activity during the blank screen prior stimulus onset in each trial and *F* is fluorescence activity in the ROI during stimulus presentation in the trial. A neuropil-correction was performed with an index of 0.7 *(**Chen et al., 2013**)*

### Sparseness measure

The sparseness measure is used to quantify the peakedness of response distribution. The sparseness measures based on the cosine distance between the response vector and the all-one vector (Rolls and Tovee, 1995; Vinje and Gallant, 2000) are popular for quantifying sparseness of responses based on spiking data. These measures however are very sensitive to slight changes in baseline level. These changes will cause a problem in our estimate of calcium signal-based sparseness which were not sensitive to spiking activities below 10 Hz (Li et al., 2017). In addition, a number of image filtering and corrections applied will also lead to an uncertain baseline in image data, which cannot even be recovered by de-convolution techniques. Thus the absolute sparseness with the Rolls-Tovee/Vinje-Gallant measure varied considerably from one study to another

An original motivation of sparseness measure is to characterize the peakedness of the distribution of neuronal population responses or the stimulus tuning curve (***Willmore et al., 2011***). In accord with this intuition, the half-height bandwidth of a tuning curve is also a simple and intuitive measure for the sparseness of neuronal responses and has been used in other studies (Rust and DiCarlo, 2012). This measure is not sensitive to low level activities or baseline fluctuations in imaging.

### Stability and reliability of the neuronal measurements

For each single neuron, we examined whether the sparse strong responses (Δ*F/F0* > 50% max) observed across the 2,250 stimuli were reliable across trials by performing the following ROC analysis (Quiroga et al., 2005): we set all the stimuli that produced mean responses greater than 50% of the observed maximum mean peak of the cell to be in the ON class, and all other stimuli to be in the OFF class. We computed the ROC for classifying the ON class against the OFF class based on the response of each single trial. If the responses above the half-maximum are stable across all trials, then the AUC (the area under the ROC curve) would be close to 1.0 as the ON and OFF classes are readily discriminable. The null hypothesis is that sparse strong responses are spurious single trial epileptic responses, thus not repeatable across trials. To test this hypothesis, we shuffled all the responses against the stimulus labels, and recomputed the mean responses for all the stimuli across the trials. We performed 1000 shuffles. We found that most of the shuffled cases have much lower average peak responses because of the mismatch of the rigorous sparse responses across trials, suggesting the reliability of the sparse responses in the original data. To make an even stricter and more fair comparison with the original data on ROC terms, for each shuffle, we recomputed the maximum responses, and used the half of this mean maximum as threshold to sort the stimuli into ON and OFF classes and repeated the ROC analysis to obtain the AUC for this shuffle. The probability of the null hypothesis is the percentage of the time that the AUCs of the 1000 shuffles reach the AUC of the original data. With this ROC analysis, we found > 96% neurons with the probability of the null hypothesis p<0.01 (Figure 1-Figure supplements 2).

### Decoding Analysis

We used a nearest centroid classifier to discriminate the 2,250 images based on the population responses in each trial. Since each image was tested 3 times, the nearest centroid classifier was trained based on two trials for all images and tested on the hold-out trials. We repeated the procedure for each trial, performing 3-fold cross-validations.

For each monkey, we constructed neural response matrices (with dimension 2250 × 1225 for monkey A, and 2250 × 982 for monkey B) for three trials X^(1)^, X^(2)^, and X^(3)^, that store the neural responses to all images in each trial as rows in that matrix. We trained and tested nearest-centroid classifiers via a three-fold cross-validation procedure across trials in a 2250-way image decoding task. Specifically, for trial t, during training, we computed the centroids of the other two trials C^(t)^ (if t=1, C^(1)^= (X^(2)^ + X^(3)^)/2, if t=2, C^(2)^= (X^(1)^ + X^(3)^)/2, etc.) and stored C^(t)^ in the classifier; during testing, given some row k of X^(t)^, which is the population neural response vector to image k in trial t, the (trained) classifier computed the Euclidean distances between row k of X^(t)^ and every row of C^(t)^. The model outputted the index (1,2,…,2249,2250) of the row in C^(t)^ that gives the smallest distance. The correct output is k and all other outputs are incorrect. The average decoding accuracy for this trial is defined as the percentage of correct outputs over all rows of X^(t)^. We repeated the above procedure for each trial and reported the average of three (average) decoding accuracies.

In our experiments, we first set the X^(t)^’s defined above to be the original recorded neural responses and computed the decoding accuracies for both monkeys. We refer to the accuracies obtained from original neural data as “achievable decoding accuracies”. Later, to evaluate the amount of information in the strong sparse portions of the neural data, we set X^(t)^’s to be thresholded versions of the original data. We tried two classes of thresholding methods: “top only” (red in Figure 2) and “top excluded” (blue in Figure 2). In “top only”, we only kept the largest p% of the responses across images and trials in the thresholded version and made the smaller (100 - p)% of the responses to be zero. In “top excluded”, which is complementary to “top only”, we set the largest p% of the responses to be zero and kept the smaller (100-p)% of the responses. For both “top only” and “top excluded”, we evaluated decoding accuracies at the following percentages (p’s) (crosses in Figure 2): 0, 0.1, 0.2, 0.3, 0.4, 0.5, 0.6, 0.7, 0.8, 0.9, 1, 2, 3, 4, 5, 6, 7, 8, 9, 10, 20, 30, 40, 50, 60, 70, 80, 90, and 99.

## ACKNOWLEDGMENTS

We are grateful to many colleagues for their insightful discussion and generous help on this paper. We thank Wenbiao Gan for the early provision of AAV-GCaMP5; and Peking University Laboratory Animal Center for excellent animal care. We acknowledge the Janelia Farm program for providing the GCaMP5-G construct, specifically Loren L. Looger, Jasper Akerboom, Douglas S. Kim, and the Genetically Encoded Calcium Indicator (GECI) project at Janelia Farm Research Campus Howard Hughes Medical Institute. This work was supported by the National Natural Science Foundation of China No. 31730109, the National Science Foundation of China Outstanding Young Researcher Award 30525016, a project 985 grant of Peking University, grants from the National Basic Research Program of China (2017YFA0105201), Beijing Municipal Commission of Science and Technology under contract No. Z151100000915070, NIH 1R01EY022247 and NSF CISE 1320651 and IARPA D16PC00007 of the U.S.A.

## AUTHOR CONTRIBUTIONS

S.T. conceived the project and designed the experiments. S.T. performed experiments with assistance from M.L., F.L., and H.J. on preparations. S.T. and T.L. analyzed the data and wrote the paper. Y.Z and Z. L. performed the decoding analysis.

---

**Figure 1 - Figure supplement 1.**
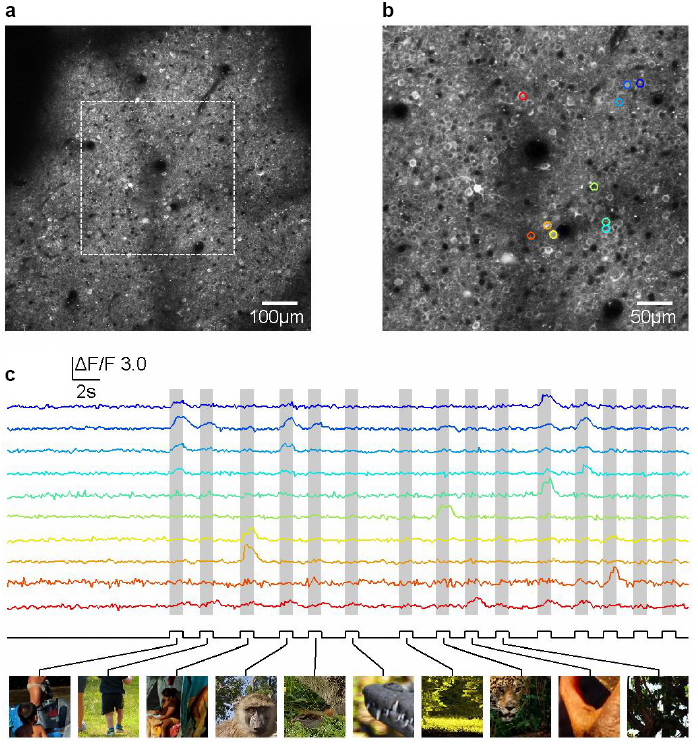
Two-photon calcium imaging in awake macaque monitoring the neuronal activity in V1 layer 2/3 evoked by natural stimuli. (a and b) Two-photon images of neurons expressing GCaMP5 at zoom 1X, 2X respectively. (c) Natural stimuli evoked robust neural activity probed by calcium indicator GCaMP5.

**Figure 1 - Figure supplement 2.**
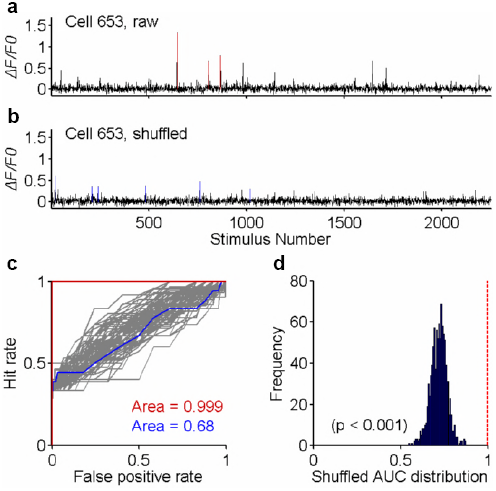
Reliability analysis of neuronal responses. (a) The tuning curve of neuron æ653’s responses (averaged across trials) to the 2250 stimuli. (b) The tuning curve computed from one random shuffle across all trials. (c) The ROC of original trials (red curve) against those of 99 shuffled trials (gray). The AUC of the original data is 0.999. The AUC of the example in (b), colored in blue, is 0.68. (d) The distribution of the AUC’s of all 1000 shuffled cases. The probability, that the shuffled AUC can reach the raw data’s AUC, is less than 0.001 (p < 0.001) for neuron æ653.

